# Guide scaffolding improves phylogenetic accuracy and enables integration across 16S amplicon regions

**DOI:** 10.64898/2026.01.05.697767

**Authors:** Holly K. Arnold, Emma Little, Alexandria Hunt, Thomas J. Sharpton

## Abstract

The growing magnitude and scale of microbiome studies now support application of meta-analytical frameworks, however the challenge of integrating microbiome sequence data across distinct studies into a unified phylogeny remains a critical barrier to progress. The 16S rDNA gene has been key to sequence-based phylogenetic microbial community analyses for over 30 years in applications ranging from human health to agricultural production efficiency. Because the full length 16S gene (∼1500 base pairs) is longer than what short-read sequencing technologies can typically capture, researchers must target a portion of the gene, or variable region, typically ∼250 base pairs in length. Using a simulation-based approach, we show that full-length, phylogenetically diverse 16S guide sequences can serve as a scaffold to improve topological accuracy of phylogenetic trees constructed from any variable region. We also show that guide sequences can provide enough essential context to build accurate tree topologies, even in the case of disjoint amplicons, allowing for integration of microbial datasets across variable regions. We validate that our findings extend to experimental data. Our approach not only suggests that guide sequences should become a standard for building phylogenetic trees from any short-read microbial data, but also extends microbial meta-analytical methods by allowing phylogenetic integration across cohorts which have used different variable regions to assess the microbiome’s association with ecosystem services. We provide an easily usable application of our method, *phyloguidesR*, as an open-sourced tool to enable any researcher to apply and use guide sequences easily in their microbial community of interest.

**IMPORTANCE:** Integrating data across independent studies can uncover robust biological patterns that strengthen inference and support more generalizable conclusions. In microbiome research, such integration is constrained by the heterogeneity of how microbial communities are sequenced. May studies target different regions of the 16S rDNA gene, producing datasets that lack a shared phylogenetic framework, and are difficult to analyze together without discarding information. As a result, comparisons across studies often rely on coarse conglomeration of taxonomic summaries or exclude entire datasets to integrate results. Here, we introduce a phylogenetic scaffolding framework that produces more accurate phylogenetic trees and demonstrate that it can unify even non-overlapping 16S amplicon datasets. Such integration has the power to reveal generalizable patterns that are invisible within individual microbiome studies, where noise and region-specific amplification have long obscured biological signals. In doing so, guides set the stage for a more powerful and integrative microbiome science at scale.

## INTRODUCTION

From optimization of human health to modulating disease, bioremediation technologies to clean energy systems, and soil science to improving our sustainable food supply, we find ourselves continually turning to one of the most ubiquitous, genomically innovative, and tiny organisms for further study – microbes. The growing magnitude and scale of microbiome studies now support application of meta-analytical frameworks (e.g. ^1–6^). Meta-analyses have been successful in many areas of molecular epidemiology, especially when rigorous integration of multiple cohorts enables analysis at a scale sufficient to identify reproducible associations while allowing for rigorous statistical control of cohort-specific confounding effects^1,7^. Meta-analyses improve power to detect associations with smaller effect sizes, as well as enable the opportunity to detect heterogeneity or comparable signals across cohorts^7^. Recent work now enable population-scale meta-analysis of microbiome data to identify consistent taxa which associate with ecosystem functions (e.g. health, disease) while appropriately controlling for confounding population structure within microbiome community data^1,8–10^. Despite these advances in microbial meta-analytical techniques, these methods commonly collapse microbial sequences into discrete taxonomic bins, and perform statistical analyses on the resulting counts, thereby discarding the rich phylogenetic information inherently contained in the sequencing data^1,8,9^.

Arguably one of the most sequenced genes on earth – the 16S ribosomal RNA (rRNA) gene – has enabled molecular profiling of microbial communities for over 30 years, and is used as a gold standard marker gene to reconstruct microbial phylogenies due to its relatively slower rates of evolution, and comparatively lower rates of horizontal gene transfer^11,12^. Microbial phylogenetics provide a richer source of information as compared to taxonomic labels alone: they capture the degree of similarity or divergence between taxa, rather than grouping microbes into coarse and sometimes arbitrary taxonomic bins. Microbial phylogenetics are also commonly used for microbial community analyses (e.g. Unifrac)^13,14^.

Importantly, microbial trait evolution is more tightly linked to phylogeny as compared to taxonomic labels, and thus many microbial traits of interest (e.g. metabolic pathways, antibiotic resistance, virulence factors) may be present in some lineages within a taxonomic label but not others^15,16^. When microbes with differing trait states are conglomerated under the same taxonomic label, key signal of how microbial traits distribute across the system, may be lost^17,18^. Additional limitations of taxonomic labels include high error rates of assigning taxonomies to short-read sequence data^19^, identical taxonomic labels with multiple ancestors (i.e. polyphyletic clades^20^), and unequal phylogenetic biodiversity contained within a taxonomic label^21^. These limitations highlight the need for phylogenetically informed methods that better preserve evolutionary signal in microbiome meta-analyses.

Limitations of taxonomic labels motivated the development of label-independent methods, which bypass taxonomic bins and instead apply statistical analyses directly to microbial sequence counts themselves. For example the burgeoning field of ecophylogenetics^17^ allows for integration of microbial phylogeny directly with microbial community distributions, eliminating the need to discard the rich information contained in microbial sequencing data prior to analyses, and does not rely on taxonomic labels at all. In this way, microbial clades – at any taxonomic depth – can be statistically associated with the ecosystem covariates of interest, enabling identification of related groups of microbes that are more prevalent than expected by chance given their position in the phylogeny and their distribution across samples (e.g. ^17,18,22–24^). Only after identifying associations with the ecosystem covariate of interest are taxonomic labels mapped onto groups of related clades, allowing researchers to describe “evolutionary hotspots” within or between taxonomic labels with particular ecosystem functions. As a result, ecophylogenetic approaches offer a powerful path to optimize interpretability and generalizability of microbiome studies.

Optimal performance of ecophylogenetic methods require phylogenetic trees as input that mirror evolutionary relationships among microbes. In practice, this presents some challenges. Most microbial datasets are produced from “short-read” second-generation sequencing technologies, which capture only ∼250 of the ∼1,500 base pairs of the 16S rRNA gene. Because full-length 16S sequences yield more accurate phylogenies^25,26^, methods that improve phylogenetic accuracy beyond what short-read sequences can produce are desired. In addition, the amount of microbial sequences produced per study are immense – for example some studies can have tens or even hundreds of thousands of unique microbial sequences which need to be integrated into a tree for downstream phylogenetic analyses. In practice, methods for tree construction need to be efficient, and heuristic approaches are frequently employed to make large-scale trees in a time efficient manner^27^.

In parallel with the increasing scale of short-read microbiome data suitable for meta-analysis, microbial reference databases of full-length representative 16S sequences have expanded substantially^28,29^. Several reference databases exist, but there is substantial effort to maintain and curate a set of up-to-date, quality-controlled rRNA genes which are both manually curated, and in which taxonomies have been checked and controlled for inconsistencies^30^. In particular ecosystem-specific contexts, such as the vertebrate gut, it is likely that most biodiversity has been described, with very close relatives already present in reference databases. It is also reasonable to assume that the amount of data in representative taxonomic databases – as well as the abundance of microbial short-read data – will continue to increase.

Our objective was to leverage the expanding availability of full-length curated 16S sequences, together with microbial short-read data to enhance ecophylogenetic analyses with three goals in mind: (A) improving the accuracy of tree construction within an individual cohort, (B) enabling ecophylogenetic meta-analyses across cohorts, and (C) providing an accessible, community-ready bioinformatic tool. We elaborate on each of these goals below.

First, we sought to determine if we could increase accuracy of phylogenetic trees derived from short-read data within an individual cohort. Ecophylogenetics requires as input a phylogenetic tree of microbial sequences present within the community of interest, and thus accuracy of ecophylogenetic methods depend upon the accuracy of phylogenetic tree construction itself. Because full-length 16S sequences can produce more accurate phylogenies compared to that of shorter partial sequences^31^, we hypothesized that concatenation of full-length “guide sequences” with microbial short-read data would provide more accurate phylogenetic tree construction, reasoning that guide sequences would provide critical phylogenetic context for improved accuracy.

Second, we sought to determine if incorporation of full-length 16S sequences could provide a viable framework for enabling ecophylogenetic meta-analyses across studies. A complication of microbial meta-analytics is that the 16S gene has nine different variable regions (VRs, abbreviated as V1 – V9). Researchers often choose a different part of the 16S gene to amplify, resulting in partially overlapping or disjoint short-reads across studies, and thus cannot be used to build an integrated phylogeny (Figure 1A). We hypothesized that guide sequences could enable integration of datasets across different VRs, reasoning that guides could provide enough phylogenetic context – even in the case of disjoint short-reads, to build integrated phylogenetic trees across VRs (Figure 1B).

**Figure 1:**
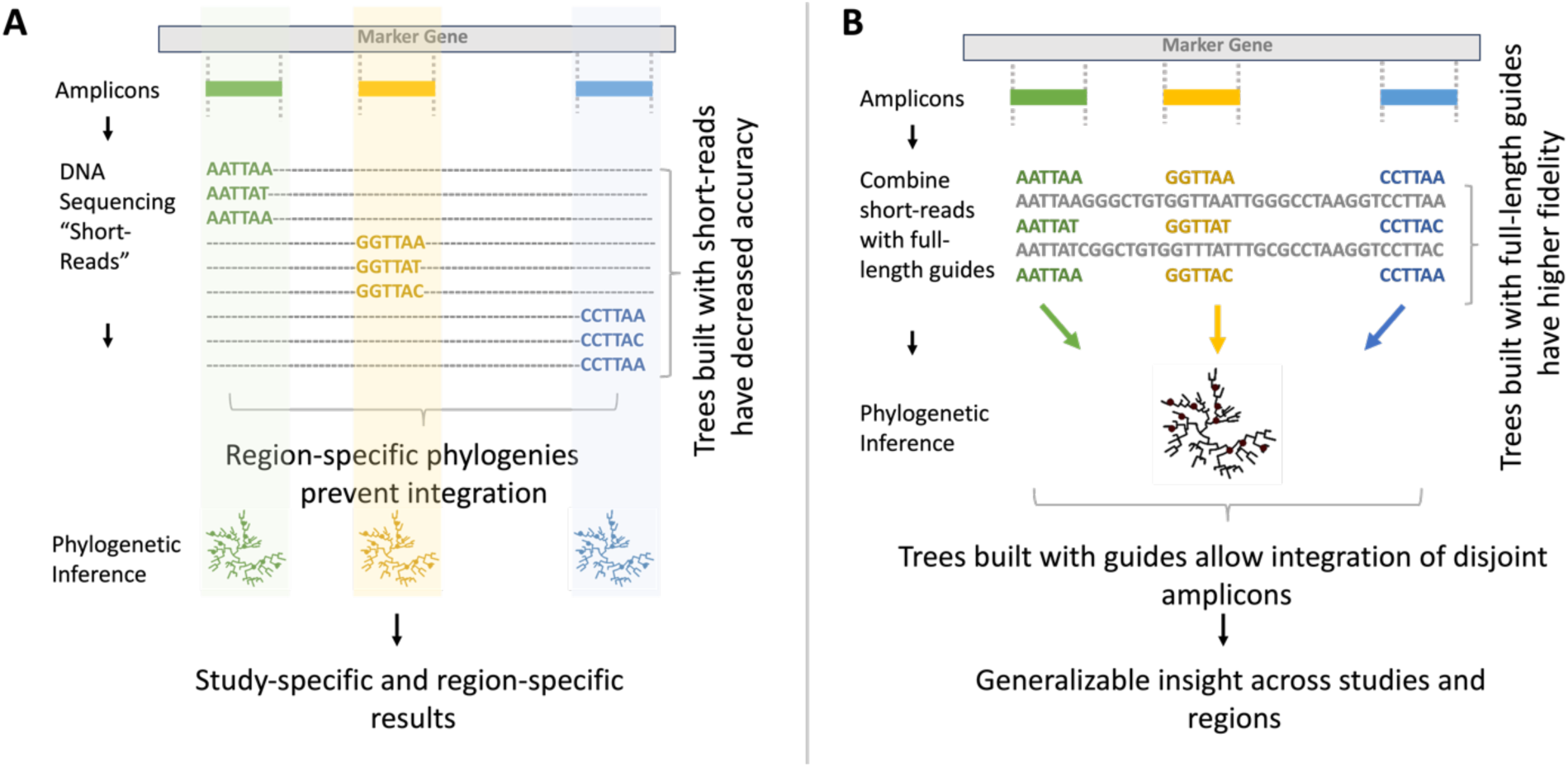
Technological limitations of high-throughput sequencing (A), and a proposed guide-sequence framework for improvement (B). **(A)** Phylogenetic trees built from short-read sequences are well known to be less accurate than trees built from full-length 16S sequences. Current high-throughput sequencing technologies generate short-reads (colored bars) that represent only a small portion of the full-length 16S gene (light gray bar). Researchers must therefore target one of nine different variable regions along the 16S gene, and phylogenetic inference is then performed on short-reads. Because different short-reads may only partially overlap, or not overlap at all, region-specific sequencing prevents integration across studies, limiting generalizability of microbiome findings for standard meta-analytic approaches. **(B)** Here, we use a framework which leverages full-length guide sequences (gray text) to scaffold short-reads within their appropriate phylogenetic context, allowing for integration of short-reads across variable regions.

Last, we sought to enable other researchers to build guided short-read 16S phylogenies. To do so, we produce an R package *PhyloguidesR* to enable researchers to apply the method to their own data with ease.

## RESULTS

To achieve our goals, we utilized a curated, high-quality, phylogenetically diverse set of full-length 16S rDNA guide sequences^30,32^ (Figure 2a), simulated microbial short-reads (Figure 2b), compared tree reconstructions with or without guides (Figure 2c), and then assessed the impact of guide-use on the accuracy of topological tree construction (Figure 2d). We evaluated six different hypotheses (H1 – H6) using the same framework:

- **H1** Guides improve phylogenetic topological reconstruction of a single variable region (Figure 2b S1).
- **H2** Guides improve phylogenetic topological reconstruction of a single variable region, at any short-read length (Figure 2b S2).
- **H3** Guides enable phylogenetic integration of <u>overlapping</u> variable regions (Figure 2b S3).
- **H4** Guides enable phylogenetic integration of <u>disjoint</u> variable regions (Figure 2b S4).
- **H5** Guides improve phylogenetic reconstruction when integrating many diverse variable regions.
- **H6:** Guides improve phylogenetic reconstruction using real experimental microbial data.

**Figure 2.**
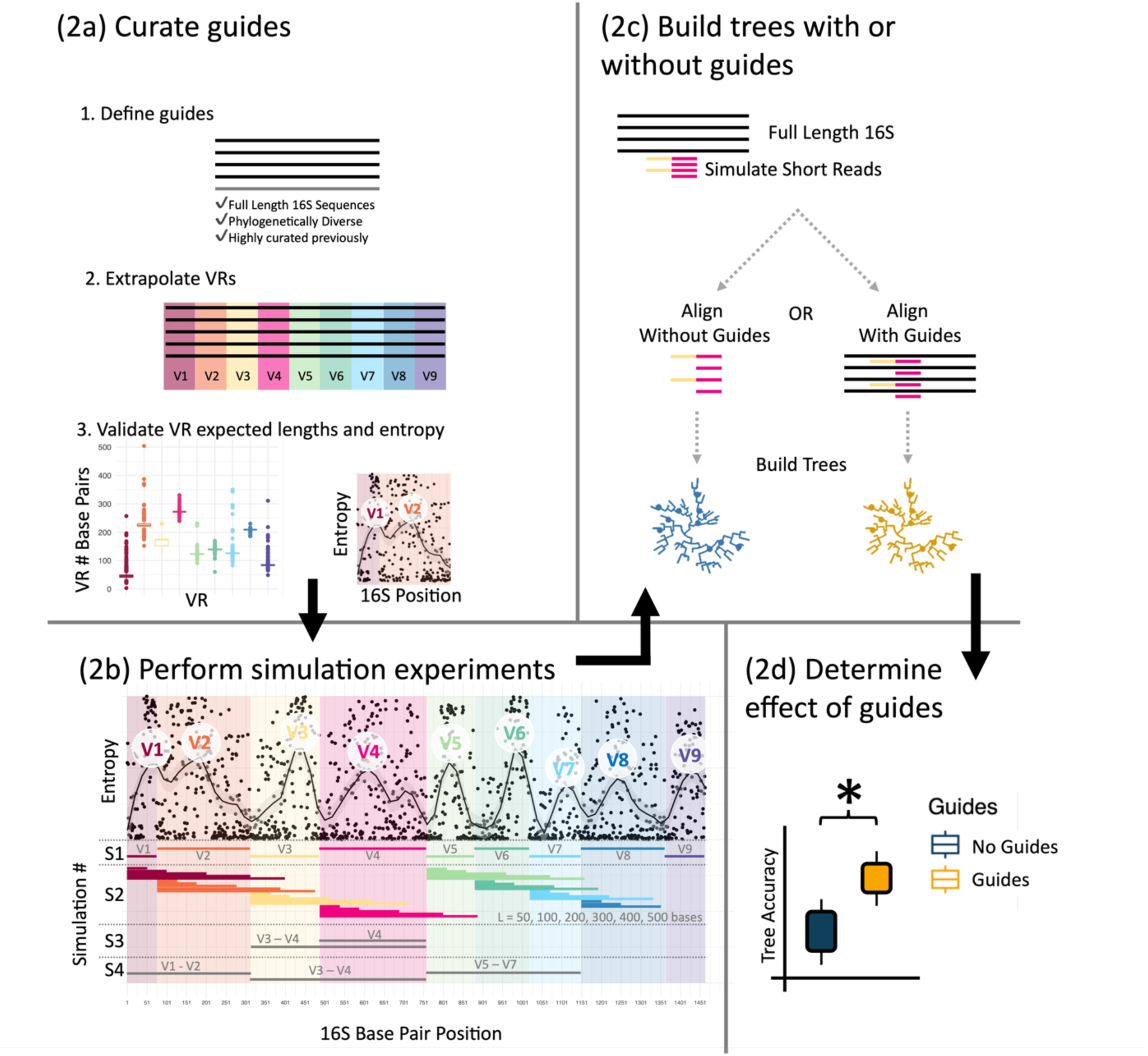
Experimental Overview: (2a) Guide sequences were derived from a curated, phylogenetically diverse set of full-length 16S sequences. Variable regions were extrapolated across guides, and then validated against expected lengths and sequence diversity (i.e. entropy). **(2b)** Short-read sequences were simulated from guide sequences in different ways, each designed to address a different hypothesis (H1 – H4). The *x-axis* shows the base pair position along the full-length 16S gene, and each simulation is designated by numbers S1 – S4 on the left side of the graph, where colored bars indicate the different short-reads produced for each simulation. **S1** generated short-reads from each variable region. **S2** generated short-reads of varying lengths (50, 100, 200, 300, 400 and 500 base-pairs) from each variable region. **S3** generated short-reads from two <u>overlapping</u> variable regions (V3V4 and V4). **S4** generated short-reads from three <u>non-overlapping</u> variable regions (V1V2, V3V4, and V5V7). The scatter plot shows how simulations align with expected sequence variation across the 16S gene, with entropy (*y-axis*) plotted by base pair position (*x-axis*). The black line shows smoothed entropy along the 16S gene. **(2c)** Next, short-reads were used alone or combined with guide sequences for phylogenetic inference. **(2d)** Finally, we assessed the impact of guide use on the accuracy of tree construction.

### Assigned VRs within guide sequences match expected patterns

To set the stage, we first set about the task of selecting a suite of full-length phylogenetically diverse 16S guide sequences (*n = 6,846*) curated previously^30,32^ (Figure 2a.1; See methods). After extrapolating VR regions across guide sequences (Figure 2a.2) we sought to validate that the regions deduced were consistent with prior work. We found guide sequence average VR length was strongly positively correlated with previously reported VR lengths^33^ (*lm; R*^2^ *= 0.946; p << 0.0001*; Figure 2a.3; Supplemental Figure 1), that per-base-pair entropy scores matched previously described sequence diversity patterns^34^ (Figure 2b; Supplemental Figure 2), and that secondary structures important for rRNA 16S functionality mapped to expected VRs (Supplemental Figure 3).

### Hypothesis 1: Guide sequences significantly improve topological reconstruction accuracy of trees across all VRs (Figure 2bS1)

It is well established that full-length 16S sequences produce more accurate phylogenetic inference as compared to short-reads alone^25,26^. We reasoned that addition of curated full-length 16S sequences alongside short-read data could provide additional phylogenetic context to improve phylogenetic inference. To test our hypothesis, we simulated microbial short-reads from each VR (V1 – V9; Figure 2bS1). For each VR, we performed phylogenetic inference using either short-reads alone or short-reads concatenated with full-length guides (Figure 2c). To assess the accuracy of inferred phylogenies, we compare short-read trees to phylogenetic trees derived from full-length 16S sequences. For the purposes of this analysis, we assume that short-read trees with higher similarity to gold standard full-length controls more closely approximate the underlying phylogenetic “truth”. Hereafter we refer to the guided group with a “+” after the VR region that was used for the simulation, and a “-” for the guideless controls. For example “V4+” would indicate short-reads simulated from the V4 region and aligned with guides, while “V4-”would indicate no guides were utilized for phylogenetic inference.

### Global tree topology is improved with guide use across all VRs

We utilized Robinson-Folds (RF) distance as a method to calculate overall distance between phylogenetic trees^35^. Guides markedly improved global tree topological accuracy regardless of VR (Figure 3a; W = 10,000; Normalized RF Distance; Wilcoxon Rank Sum Test; p << 0.0001). In the guided group, V2+ performed significantly better than V4+ (Wilcoxon Rank Sum Test; p << 0.0001), but in the guideless group, V4-performed slightly better than V2-(Wilcoxon Rank Sum Test; p << 0.0001), implying that guides improve phylogenetic inference in a VR-dependent manner. Despite the gains in accuracy using guides, no guided VR was able to reach the accuracy of full-length controls (Wilcoxon Rank Sum Test; p << 0.0001), indicating that guides provide valuable contextual information for phylogenetic inference, but are not able to match the full-length 16S gene for accuracy.

**Figure 3:**
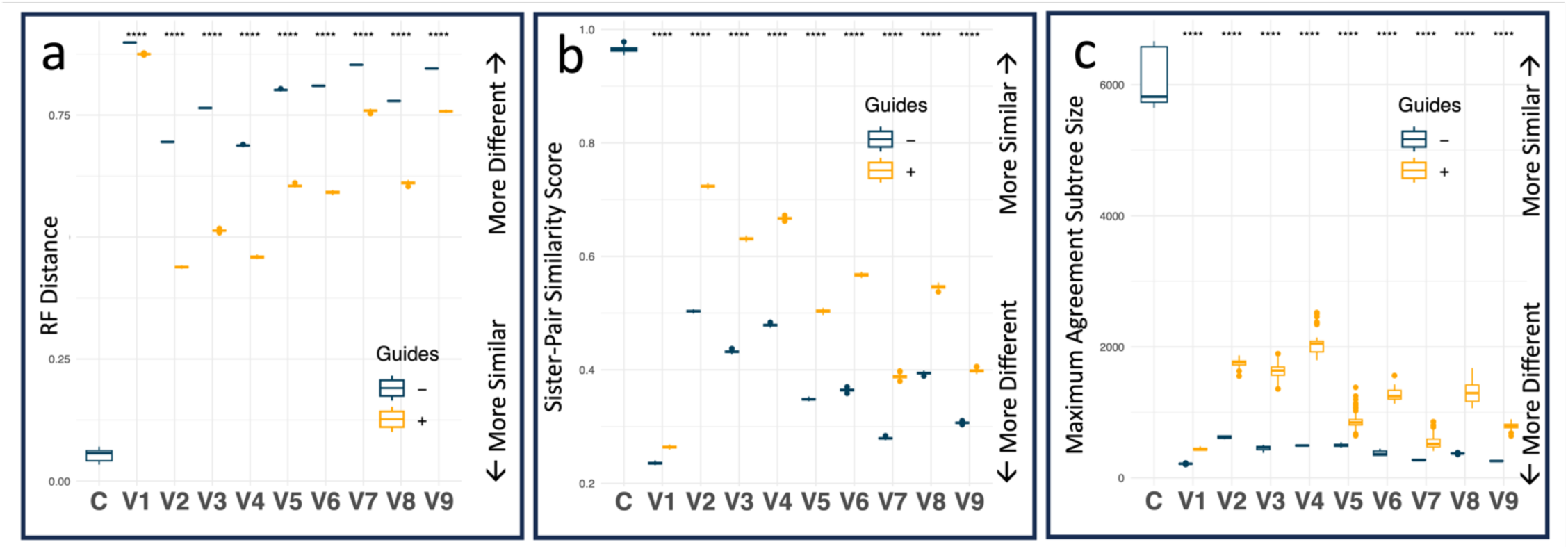
Global and local tree topology is improved with guide use across all VRs. **(a)** Plot shows normalized Robinson Fold (RF) distance of short-read phylogenetic trees (V1 – V9) as compared to gold standard controls (0 = identical trees, 1 = maximally different trees). Comparison of two sets of independently generated full-length trees serves as a control (C). **(b)** Plot shows the number of shared sister pairs (*y-axis*) as a function of VR and guided group (*x-axis*). (1 = 100% identical sister pairs, 0 = no shared sister pairs). **(c)** Maximum agreement subtree size (*y-axis*) as a function of VR and guide group (*x-axis*). Statistical significance notated as **** indicates Wilcoxon Rank Sum Test p ≤ 0.0001.

The RF^35^ tree distance metric is arguably the most widely cited metric for topological tree comparison^36,37^, but has been criticized for becoming rapidly saturated^37,38^. To address this limitation, we also tested our findings across a battery of additional tree distance metrics - developed for cases when subtree structures are highly similar, but not exactly identical (Supplemental Table 1) – including generalized RF distances (Jaccard Robinson-Foulds^36^ distance, Nye^39^ distance, matching split distance^40^), information theoretic generalized RF distances (phylogenetic information distance^37^, mutual clustering distance^37^), as well as metrics which consider global pairwise distances (path distance^41^), or quartet substructure (quartet distance^42^). Overall, we found that guide use significantly improved topological tree accuracy across all VRs, and that this result was robust, no matter which global distance metric was considered with some minor exceptions (see Supplemental Figure 4 for further discussion).

### Guide sequences strongly enhance fine-scale topological accuracy across all VRs

While the aforementioned metrics capture overall similarity between two trees, they may not accurately depict how well recently diverged taxa are recovered. For example, a short-read tree may differ markedly based on global topology because deep nodes are not resolved, yet still have accurately resolved recently diverged clades. In the case of the microbial researcher, they may often times be more interested in resolving shallow, rather than deep evolutionary relationships because many traits of interest (e.g. antibiotic resistance, metabolic specialization, pathogenic islands) may arise within recently diverged lineages. To capture how similar fine-scale relationships are between short-read and gold standard trees, we developed a cladal similarity metric which quantifies how often two trees group the same taxa together in clades of a given size, independent of tree branching order. We convert this to a number between 0 - 1 which can be intuitively thought of as the proportion of clades of a given size which have identical membership between the two trees (see methods for details). We calculate a cladal similarity metric for sister pairs, trios, quadruplets,…, up to decuplets (clades of size 10).

Guides markedly improved recovery of fine-scale phylogenetic relationships across all VRs (Figure **3b**; Wilcoxon Rank Sum Test; p << 0.0001). For example, in V2, guides boosted sister-pair recovery by ∼22%, capturing 72% ± 0.24 of true sister pairs as compared to only 50%± 0.17 when not using guides. A similar effect was seen for V4, where guide use allowed for recovery of ∼19% more sister pairs when using guides (𝜇_𝑉4+_ = 0.67; 𝑆𝐷_𝑉4+_ = 0.0020; 𝜇_𝑉4_ = 0.48; 𝑆𝐷_𝑉4_ = 0.0017). We extended this analysis to consider clades with not only identical sister pairs (size = 2), but identical sister triplets, quadruplets, …, up to decuplets, and found similar patterns (see supplemental Figure 5 for details), showing that guides not only improve overall global phylogenetic inference, but also rescue fine-scale relationships of shallowly diverged taxa.

Additionally, we calculated Maximum Agreement Subtrees (MAST) between short-read and gold-standard trees. MAST is defined as the largest subtree that can be obtained between both trees by deleting, but not rearranging nodes^43^. We reasoned that the MAST represents an intermediate zone of phylogenetic similarities between global (e.g. RF distance) and microscale (e.g. cladal similarity metric) topology. MAST reflects the extent to which a coherent backbone exists between two trees despite localized errors, whereas the cladal similarity metric focuses on local neighborhoods and global similarity metrics (e.g. RF) focuses on the total tree structure.

Guides substantially increased coherence between phylogenies when MAST size was considered (Figure 3c). Across every VR, guides allowed for significantly larger MASTs to be recovered accurately as compared to unguided trees (Wilcoxon Rank Sum Test; p < 0.0001). The effect was most striking for V4 where guides increased MAST size by over threefold – from 494 ± 5.5 tips in unguided trees to 2,041 ± 161 tips with guides, an improvement of 1,548 tips of shared tree structure. As expected, control full-length trees produced MASTs that were nearly identical to total tree size (6,118 ± 424; 𝑁_𝑇𝑅𝐸𝐸_ = 6,846). Taken together, simulations support that guide use improves phylogenetic inference, and that this effect is true for any VR of consideration.

### Hypothesis 2: Guide sequences improve topological accuracy across VRs and read-lengths (**Figure 2b**S2)

Short-read datasets vary in length. We sought to determine whether a critical short-read length is required for guides to yield improvements in phylogenetic inference. We hypothesized that longer read lengths would yield more accurate phylogenetic inference overall, and that the relative benefit of guides would plateau as read length increased. We reasoned that longer reads have more intrinsic phylogenetic signal, thereby reducing the additional gains provided by guide sequences. To test our hypothesis, we simulated varying read lengths (L = 50, 100, 200, 300, 400, 500 base pairs) across VRs (Figure 2bS2). For each VR and read length, we performed phylogenetic inference using either short-reads alone or short-reads concatenated with full-length guides (Figure 2c). As before, to assess the accuracy of inferred phylogenies, we compare short-read trees to phylogenetic trees derived from full-length 16S sequences.

Across all variable regions and read lengths, guides significantly improved both global (Supplemental Figure 6) and fine-scale (Supplemental Figure 7) tree accuracy (Wilcoxon Rank Sum; p << 0.0001). Accuracy improved with increasing read length most substantially near the ∼300 base pair mark, indicating that guides significantly provide phylogenetic inference accuracy gains at read-lengths routinely generated by high-throughput amplicon sequencing. This highlights their practical utility for maximizing phylogenetic accuracy in short-read datasets. As expected, guides resulted in decreasing accuracy gains as read length increased.

Regression analyses show that read length, VR, and guide use interact to influence accurate phylogenetic inference, with guides consistently enhancing both global topology (Beta Regression; logit link; Pseudo R2 = 0.80; p << 0.0001; see Supplemental Table 2 and discussion) and fine-scale sister pair recovery (Beta Regression; logit link; Psuedo R2 = 0.78; p << 0.0001; see Supplemental Table 3 and discussion). MAST sizes were also significantly increased in all simulations when guides were used, regardless of length or VR (Supplemental Figure 8; Wilcoxon Rank Sum Test; p << 0.0001). Strongest effects were noted in V2+ and V4+. Taken together, our simulation results indicate that common short-read lengths found in experimental data benefit from guide scaffolding when performing phylogenetic inference.

### Hypothesis 3: Guides enable phylogenetic integration of overlapping VRs (Figure 2bS3), outperforming all guideless methods

The growing magnitude and scale of microbiome studies now support application of meta-analytical frameworks to combine microbial 16S data across many studies, however a challenge of combining microbiome sequences derived from distinct VRs remains a critical barrier to progress. As a result, researchers often exclude studies which have not sequenced identical VRs, resulting in data loss, and limiting the generalizability of findings across studies. We hypothesized that guide sequences could enable phylogenetic integration of 16S data derived from distinct but overlapping VRs (e.g. V4 and V3V4) with reasonable accuracy. Here, we define reasonable accuracy as phylogenetic inference performance that is at least as accurate as that obtained using current state-of-the art methods (i.e. unguided short-read data from a single VR region alone). To test our hypothesis, we simulated short reads from one VR alone (V4 or V3V4), or randomly mixed V4 and V3V4 reads into a single group (Mix).

We found that the mixed guided group (Mix+) consistently outperformed all guideless groups (V4-, V3V4-, Mix-) across global topology (i.e. RF), recovery of fine-scale relationships (i.e. sister pairs), and phylogenetic backbone (i.e. MAST) (Figure 4, Wilcoxon Rank Sum Test; q < 0.05). Notably, the guided Mix+ group surpassed the guided V4+ group in RF distance as well as in recovery of sister pairs, likely reflecting gains from both overall longer read length within the group and scaffold integration. These results show that guides not only improve phylogenetic reconstructions of individual VRs, but also provide a robust framework for integration of overlapping VRs across studies.

**Figure 4:**
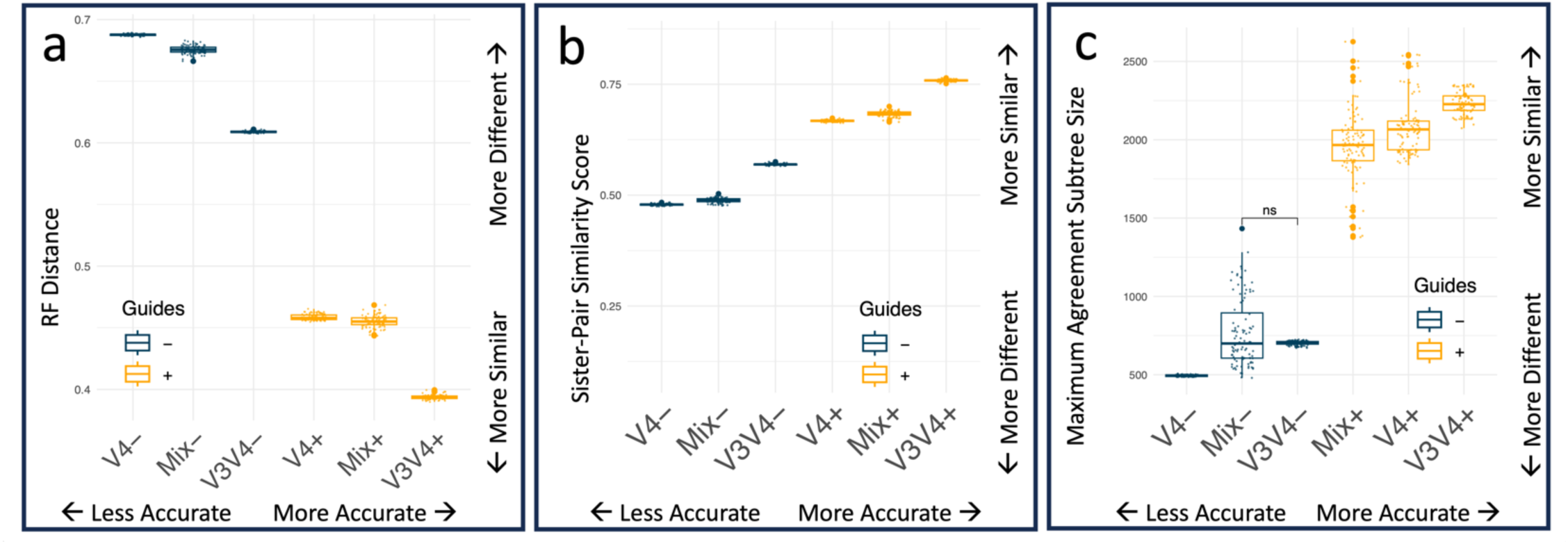
Guides enable phylogenetic integration of overlapping VRs. Plots show the accuracy of phylogenetic trees built with (+) or without (-) guides from simulated short reads derived from V4, V3V4, or a mix of V4 and V3V4 (denoted as “Mix”). RF Distance (a), sister pair recovery (b), and MAST size (c) are displayed (*y-axis*) as a function of simulation group (*x-axis*). Boxes are ordered by increasing median accuracy. Statistical significance of all pairwise combinations between groups were calculated via Wilcoxon Rank Sum test, then corrected for multiple hypothesis testing. Unless otherwise indicated (“ns” = non-significant), all pairwise comparisons were significantly different (q <0.05).

Robustness analyses confirmed these findings: across eight global topological metrics, the guided Mix+ group significantly exceeded the best unguided group, and even surpassed accuracy of the V4+ in three of eight of the metrics (Supplemental Figure 9). Analysis of the robustness of local accuracy results yielded similar conclusions – the Mix+ group consistently improved recovery compared to all unguided groups, regardless of what clade size was considered (Supplemental Figure 10). Taken together, our simulations show that use of guide sequences can enable phylogenetic integration of overlapping VRs with reasonable accuracy, and can improve phylogenetic inference of single and mixed overlapping regions beyond the current state-of-the art.

### Hypothesis 4: Guides enable phylogenetic integration of disjoint VRs (Figure 2bS4), outperforming unguided methods

Given that guides provided sufficient phylogenetic context to integrate overlapping short-reads from different VRs, we next assessed whether guides could enable integration of disjoint, non-overlapping VRs with reasonable accuracy. Here, we define reasonable accuracy as phylogenetic inference performance that is at least as accurate as that obtained using current state-of-the-art methods – namely using unguided short-read data from a single VR region alone. Although disjoint VRs are, by definition, impossible to align directly, we hypothesized that guide sequences could provide the necessary scaffolding to enable their phylogenetic integration, thereby improving accuracy relative to unguided tress. To test this hypothesis, we evaluated short-reads derived from three non-overlapping regions (V1V2, V3V4, and V5V7), and analyzed them individually with and without guides. We also combined them into a mixed dataset (Mix) – where the unguided Mix-group served as a negative control.

Guided integration of disjoint VRs (Mix+) consistently outperformed all unguided controls across global topology (RF distance), local accuracy (sister-pair recovery), and MAST size (Figure 5, Wilcoxon Rank Sum Test; *q < 0.05*). As expected, the Mix-group performed poorly, demonstrating the clear importance of guides for integrating disjoint regions. By contrast, the Mix+ group achieved accuracies comparable to, and in some cases exceeding, guided single-region controls, depending on the metric considered. For example, the Mix+ group outperformed or equaled guided controls in certain global metrics (MSD, PID, Path and quartet distances; Supplemental Figure 11) as well as at some local clade sizes (8 and 10; Supplemental Figure 12). These findings demonstrate that guide scaffolding retains high fidelity even when combining non-overlapping regions. Together these results demonstrate that guide sequences extend the current state-of-the-art by enabling accurate phylogenetic reconstruction from even disjoint VRs, something not possible with unguided approaches.

**Figure 5.**
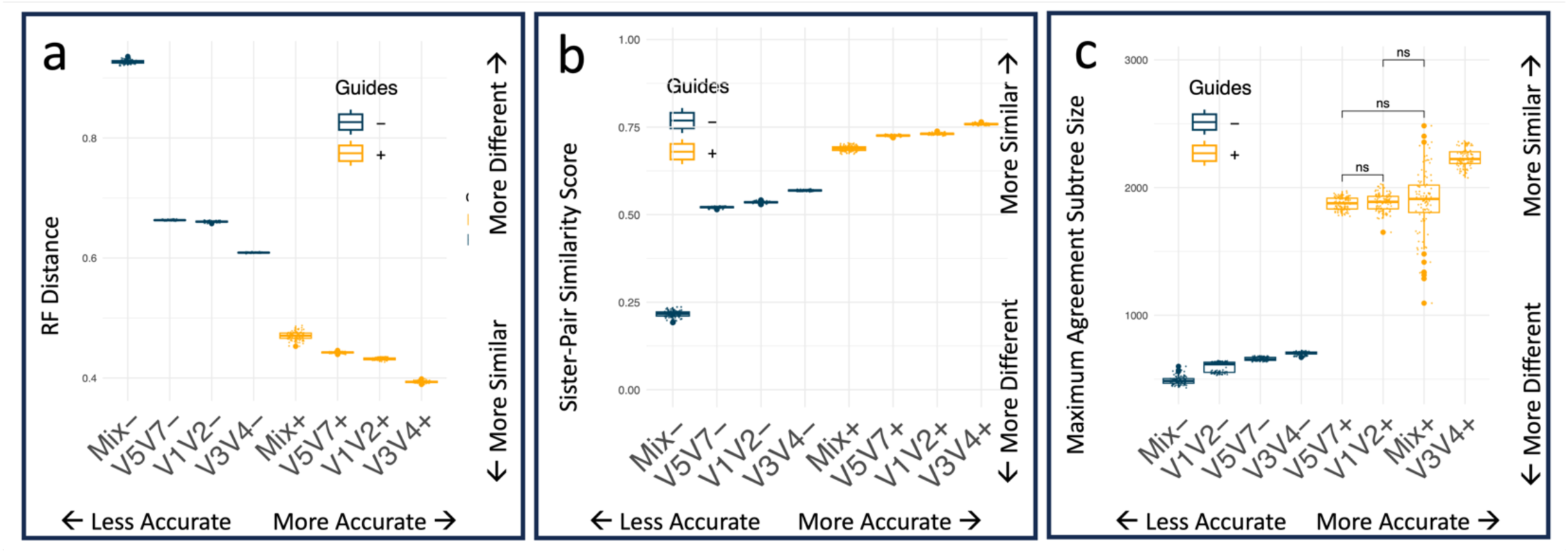
Guides enable phylogenetic integration of disjoint VRs. Plots show the accuracy of phylogenetic trees built with (+) or without (-) guides from simulated short-reads derived from single-region controls (V1V2, V3V4, and V5V7) as well a mix of all single-regions (Mix). RF distance (a), sister pair recovery (b), and MAST (c) are displayed (*y-axis*) as a function of simulation group (*x-axis*). Boxes are ordered in increasing median topological accuracy. Statistical significance of all pairwise combinations between groups were calculated via Wilcoxon Rank Sum test, then corrected for multiple hypothesis testing. Unless otherwise indicated (“ns” = non-significant), all pairwise comparisons were significantly different (*q <0.05*).

### Hypothesis 5: Guides enable phylogenetic integration of diverse VRs

Meta-analytical approaches combining multiple microbiome studies may need to integrate a diverse suite of VRs, not just two or three. We hypothesized that guides would enable phylogenetic integration of a diverse mix of overlapping VRs, improving accuracy relative to unguided trees. To test our hypothesis, we evaluated the scenario of combining a highly diverse set of short-reads from six overlapping and non-overlapping regions, and analyzed them individually with and without guides (Supplemental Figure 13, 14, and 15). Similarly to prior results, the Mix+ group performed significantly better than guideless groups in global distance (RF Distance), local distance (sister pairs), and MAST size (Wilcoxon Rank Sum; FDR; q < 0.05). Robustness analysis showed that even when considering a more diverse mix of VRs, that use of guides allowed the for the Mix+ group to either outperform all guideless groups, or perform at least as good as a control single-region guided group. Taken together, our simulation results suggest that guides can improve upon the current state-of-the art approaches to building trees from short-read data from a single VR, and enable a platform to integrate sequences across VRs.

### Hypothesis 6: Guides improve phylogenetic inference when using real experimental short-read data

To test whether the benefits of guides observed in simulations translated to experimental data, we compared Illumina short-read amplicons with matched PacBio long-read 16S rDNA datasets generated from an identical sample set^31^. Guided trees showed markedly improved accuracy, with significantly lower RF distances than guideless trees (Wilcoxon Rank Sum; p<<<0.0001; Figure 6), and consistent gains across nearly all global metrics (Supplemental Figure 16; Wilcoxon Rank Sum, p < 0.05). We found that guide-use improved the number of sister pairs that were recovered (Wilcoxon Rank Sum; p << 0.0001), and allowed larger MAST sizes (Wilcoxon Rank Sum; p <<< 0.0001). Together, these results confirm that guide scaffolding improves both global and fine-scale phylogenetic accuracy and can extend to short-read experimental datasets.

**Figure 6.**
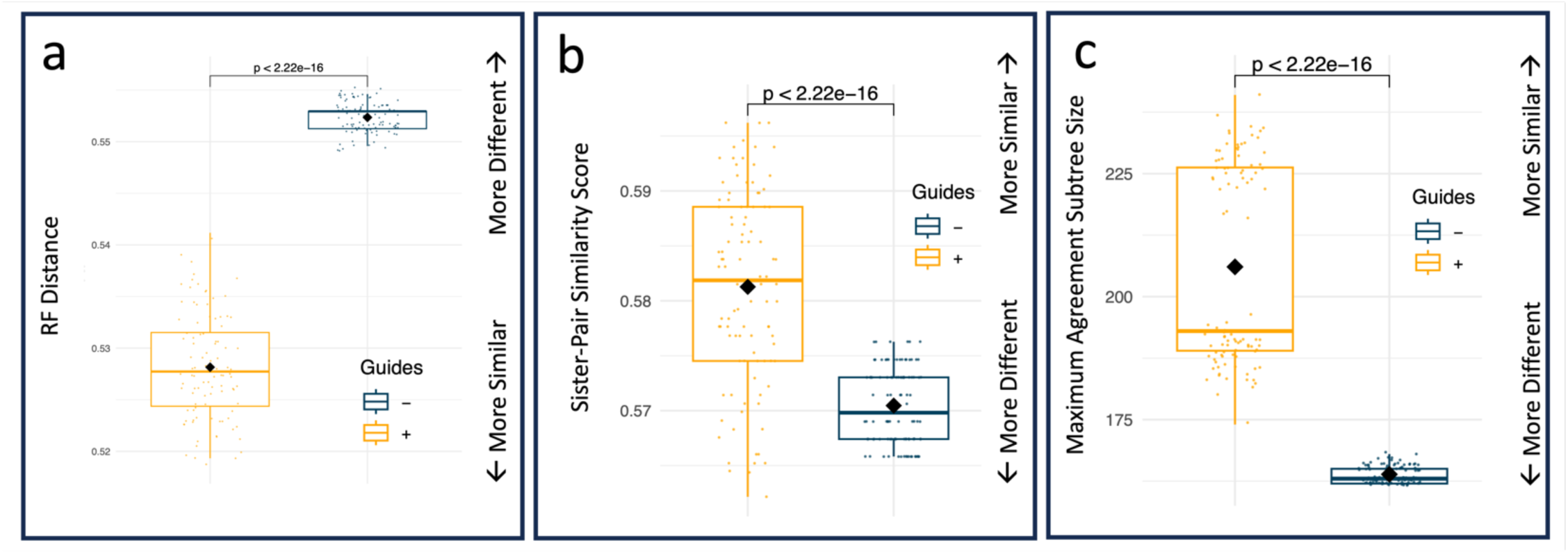
Guide sequences improve accuracy of short-read phylogenies relative to long-read trees. Trees built from Illumina short-read amplicons with and without guides were compared to PacBio full-length 16S rDNA sequences from identical samples to assess accuracy. Plots show **(a)** global topology accuracy measured by normalized RF distance (0 = identical; 1 = maximally different), **(b)** local accuracy measured by recovery of shared sister pairs (0 = none shared; 1 = all shared), and **(c)** MAST size between short and long-read trees. Statistical significance was determined via Wilcoxon Rank Sum test.

### PhyloguidesR: An accessible R package for guide-based phylogenetic reconstruction

We developed *phyloguidesR*, a lightweight and accessible R package, that enables researchers to seamlessly use guide sequences for phylogenetic inference. The package provides an intuitive workflow and wraps common bioinformatic tools (e.g. FastTree^27^, mothur^44^) in an easy to use R interface. To facilitate adoption, we provide open access to an accompanying tutorial with a demo from real 16S data^45^, allowing users to learn, then immediately have the skills to extend the framework to any short-read amplicon data and guide set. By lowering technical barriers, *phyloguidesR* makes guide-based phylogenetic integration broadly applicable to ecologists.

## DISCUSSION

Our results demonstrate that guide sequences address two major long-standing barriers in microbiome research. First, guide sequences significantly improve the accuracy of phylogenetic inference within a single VR, advancing the current state-of-the-art for phylogenetic analysis from short-read 16S data. Second, we show that guides provide sufficient additional phylogenetic context to overcome a persistent limitation in the field: they enable construction of integrated phylogenies from a heterogenous mixture of disjoint or overlapping VRs, thereby facilitating integration of microbiome results across 16S datasets.

We believe this approach has the ability to transform microbial meta-analytic methods as current approaches rely first on classifying microbial sequences to taxonomic labels, and thereby discard rich information inherently present in sequencing data. Our work sets the stage for enabling phylogenetic extension of molecular epidemiological studies which go beyond conglomeration of taxonomic labels alone. Guides sequences act as anchors for short-reads, restoring evolutionary context that would overwise be lost. By preserving evolutionary relationships among microbes, this approach will allow for conserved and divergent microbial lineages to be associated with ecosystem services which may be lost when looking across taxonomic labels alone.

Phylogenetic comparison of trees has traditionally focused on overall, or global (dis)similarity metrics between trees, however in microbiome science the most relevant signals may occur at shallow evolutionary time scales (e.g. recently acquired traits via horizontal gene transfer). In order to address this gap, we introduced a cladal similarity metric which quantifies congruence of cladal membership between trees. This new lens shows that smaller clades are recovered with higher fidelity in the presence of guides, implying that guides can enhance capture of fine scale tree topology. Guides sequences may therefore be particularly important for constructing short-read phylogenies used in recently developed ecophylogenetic methods^17,18,24,46^, that link related groups of microbes to ecosystem functions.

Our study necessarily made several simplifications and assumptions. First, our study sought to understand the impact of guide sequences for accurate phylogenetic inference independent of variation that can be introduced by alignment algorithms. Thus, we chose to focus on a reference-based alignment strategy – the Nearest Alignment Space Termination (NAST)^47^ algorithm - because this method rapidly aligns thousands of sequences with a deterministic solution. Future work should determine how alignment strategies such as de-novo multiple sequence alignment (e.g. MAFFT^48^), or other alignment approaches, influence downstream tree construction. Second, we deliberately tested the case where each short-read was paired with a guide (i.e. a 1:1 short-read-to-guide ratio), and future studies should assess how different guide-to-read ratios and phylogenetic breadth of guides affect outcomes. Third, we only evaluated guide use with a single phylogenetic inference method, because prior work^49^ has shown high concordance across algorithms. Future work should seek to determine how guide use impacts trees built from differing inference methods. Last, experimental data was limited to host-associated microbiomes, which are well represented in reference databases; testing guide performance in less-characterized environmental conditions will be critical to establish their general utility.

Overall, we address a long standing barrier in the field of microbial ecology by providing a scaffolding for integration of short-reads across VRs, thereby establishing a foundation for more powerful meta-analytical approaches to understand how microbiomes associate with ecosystems and ecosystem services. Such integration has the power to reveal generalizable patterns that are invisible within individual studies, where noise and region-specific amplification have long obscured biological signals. In so doing, guides set the stage for a more powerful and integrative microbiome science at scale.

## METHODS

### Defining the guide sequence set

Full-length 16S rDNA sequences were derived from the Species Living Tree Project^28^ (SILVA version 138.2) containing 510,495 sequences which are not more than 99% similar to one another. We chose to utilize a subset of these sequences, called “seed” sequences - previously curated for reference alignment^32^ due to their suspected ability to produce stable alignments as well as the fact that they are phylogenetically diverse. Sequences taxonomically labelled as eukaryotes (*n = 1,850*) were excluded, and only bacterial (*n = 6,714*) and archaeal (*n = 132*) sequences were considered for downstream analyses (*n = 6,846*). The seed alignment (50,000 characters in width) was filtered to remove gaps common across all sequences (5,805 characters in width) using mothur (version 1.40.4; *filter.seqs()*). Hereafter, we refer to this as the “guide alignment”. By removing gaps from the guide alignment file, a set of phylogenetically diverse full length guide sequences were derived that could be used downstream for combination with microbial short-read sequences.

### Entropy calculation of variable regions

In order to determine if our guide sequences fit reasonable expectations of conservation and variation across the defined VRs, we assessed entropy as a function of base pair position along the 16S gene. Entropy was calculated for each base position along the model *E. coli* 16S gene as 𝐻_𝑏𝑝_ = − ∑^4^ 𝑝_𝑖_𝑙𝑜𝑔_2_𝑝_𝑖_ where 𝑝_𝑖_ is the probability of finding each base 𝑖 = {𝐴, 𝐶, 𝑇, 𝐺} at each base pair position across all guide sequences. Locally estimated scatterplot smoothing (LOESS) was then used to smooth entropy values along the 16S gene. LOESS estimates fit at a particular point *x*, made by points in the neighborhood of *x*, weighted by the distance from *x*, where the neighborhood size is the proportion 𝛼 of the points. We tuned 𝛼 by considering which 𝛼 𝜖 {0.1, 0.2, . . . , 1} minimized mean squared error of predicted values of the model, and found 𝛼 = 0.1 to be best. We used the ggplot2 package (*geom_smooth(span = 0.1, method = “loess”, se = TRUE;* version 3.5.1) to visualize the smoothed function fit and data.

*Simulation of microbial short-read sequences*.

To simulate microbial short-reads, we first needed to define VR start and stop locations in the guide-alignment file. To do so, we first searched the literature for primers which are commonly used to amplify particular VRs (Supplemental Table 4). We next identified a full-length 16S rDNA sequence within our guide sequence set which was an *Escherichia coli strain* AE005174^50^ (AE005174.Esch1125) to serve as a “model species” to determine primer start location for each VR. We then identified start and stop locations of each VR via exact matching of primers to the model *E. coli* sequence. We then extrapolated the start and stop locations of each VR across all our guide sequences using our model *E. coli*’s alignment to all other guide sequences. Short-reads were then simulated by removing all gaps in the alignment file between the desired start and stop location for the particular short-read. For each simulation, one short-read was simulated for every guide sequence.

### Alignment of short-reads with and without guides

For each simulation, short-read sequences were concatenated with full-length guide sequences (+guides) or used alone (-guides) for alignment. Sequences were aligned using an ‘align to seed’^30^ approach using the Nearest Alignment Space Termination^47^ (NAST) algorithm for multiple sequence alignment (mothur version 1.48.2 *align.seqs(template=seed_alignment)*)^44^. The guide alignment file was utilized as a template. We deliberately chose the NAST algorithm as a first pass to answer our questions for several reasons, and acknowledge that many other alignment strategies exist. First, the NAST alignment algorithm considers each sequence in context of a template and thus produces a stable alignment no matter what order the sequences are considered (unlike de-novo approaches), and thus allowed more precise control across simulations. Second, NAST alignment strategies are particularly suited to highly conserved genes (e.g. the 16S rDNA gene), for which it is desired to map known characteristics of reference sequences back to the query sequence in question (e.g. the VR). Third, NAST is well suited to align thousands of microbial sequences in efficient time, making the choice relevant for microbial studies aligning tens of thousands of reads.

### Phylogenetic Tree Construction

Phylogenetic trees were constructed using a generalized time reversible model of nucleotide evolution implemented in FastTree^51,52^ (*-nt -gtr -gamma*; version 2.1.11). FastTree implements an Approximately-Maximum-Likelihood algorithm, which is appropriate for large alignments, and can run up to 1,000 times faster than other non-heuristic tools^51^. We chose to only implement one tool for phylogenetic inference as prior work has displayed high topological concordance between FastTree, IQ-Tree, RaxML-NG, and PhyML^49^. If guides were used to build the phylogenetic tree, they were subsequently dropped out of the resultant tree using the *drop.tip()* function from the ape package (version 5.8.1).

### Calculation of topological similarity between microbial short reads and full length sequences

To quantify how closely short-read phylogenies capture “true” evolutionary relationships, we first constructed a set of gold standard trees using full-length 16S sequences. Because inferred phylogenetic tree topology can exhibit sensitivity to alignment sequence order in heuristic search, we employed a bootstrap-style approach, where the order of sequences in the alignment file were randomly shuffled prior to phylogenetic inference, generating 100 replicate trees per alignment. To evaluate the accuracy of short-read trees constructed with or without guide sequences, each short-read tree was compared to the corresponding full-length gold standard tree by calculating tree distances using 100 random pairings per condition.

Global tree similarity metrics were calculated for each pair of trees to capture overall similarity between short-read and gold-standard tree topologies. We assume that short-read trees which exhibit higher similarity to gold-standard trees recapitulate true evolutionary relationships to a greater extent. For each pair of trees, we calculated eight different global distance metrics (see Supplemental Table 1 for details). Robinson Foulds distance was calculated using the phangorn package (*RF.dist(normalize = TRUE)*, version 2.12.1). Jaccard Robinson Foulds (*JaccardRobinsonFoulds(normalize = FALSE)*), Nye Distance (*NyeSimilarity(similarity = FALSE, normalize = TRUE*), Matching Split Distance (*MatchingSplitDistance(normalize = FALSE)),* Phylogenetic Information Distance (*PhylogeneticInfoDistance(normalize = TRUE)*), Mutual Clustering Distance (*ClusteringInfoDistance(normalize = TRUE)*, and Path Distance (*PathDist()*), were all calculated using the modified BigTreeDist version of the TreeTools^53^ package (version 2.6.3.9001; devtools::install_github(“ms609/TreeTools”, ref = “more-leaves”)). Path distance and Quartet distance was calculated using the tqDist package^54^ (version 1.0.2 implemented in C++) and was normalized using the theoretical maximum quartet distance of the number of ways to choose four tips from all guide sequences (*N_tips = 6,846; QUARTET_MAX = choose(N_tips, 4))*.

While global metrics capture global topology between trees, they may not capture how accurate local tree topology is resolved. For example, a short-read tree may differ markedly in global topology compared to the gold standard because deep nodes are not resolved, yet still have similarly resolved recently diverged evolutionary relationships. We quantified the accuracy of recovering shallow (fine-scale) evolutionary relationships by calculating a cladal similarity metric which compares clade membership between two trees. We focused on clades of small sizes *s* ∈ *{2, 3, …, 10}*. The clade similarity score for clades of size *s* is defined as the proportion of clades with identical membership shared between two trees, and thus ranges from 0 (no clades of size *s* are shared between trees) to 1 (all clades of size *s* are shared between trees). To calculate the metric, trees were midpoint rooted, and all clades of size *s* were extracted and membership was compared between short-read and the gold-standard trees. In the event that a clade had identical membership between trees, it was counted as shared, and otherwise was counted as unshared. Note that clade membership had to match identically between trees to be counted as shared and partial matching was not considered. In addition, this metric does not take into account branching order or branch lengths between trees. This metric provides a size-specific measure of shallow topological concordance between two trees.

We also calculated Maximum Agreement Subtree (MAST) size between short-read and gold standard trees. MAST is defined as the largest subtree that can be obtained from both trees by deleting but not rearranging nodes. Thus, MAST represents an error tolerant measure of a “shared backbone” between two trees. MAST was calculated with the phangorn package (*mast()*, version 2.12.1).

### Simulation specific details

Each of the simulations described below varied in how sequences were simulated, but used the framework described above unless otherwise specified.

Hypothesis 1 sought to determine if guides improved topological accuracy across all VRs. To test this, we simulated microbial short-reads for each VR (V1 – V9). For each VR, simulated short-reads were either used alone (designated V1-, V2-, …, V9-), or combined with the curated full-length guide sequences (designated V1+, V2+, …, V9+) for phylogenetic inference (Figure 2b.S1 and 2c). For each set of short reads, we calculated aforementioned global tree distance metrics, local cladal similarity metrics, as well as MAST sizes as compared to the gold standard trees. We evaluated the effect of guides using Wilcoxon Rank Sum Tests to determine whether the inclusion of guides significantly improved tree similarity scores.

Hypothesis 2 sought to determine if guides improved topological accuracy across a range of VRs and short-read lengths. To test this, we simulated short-read sequences at varying lengths (L = 50, 100, 200, 300, 400, and 500). Short-reads were either used alone or combined with curated full-length guide sequences for phylogenetic inference (Figure 2b.S2 and Figure 2c). We only considered simulations where all short-read sequences could be simulated to the intended length. Notably, VRs toward the terminal end of the 16S gene were excluded (V6 at 500 base pairs, V7 at 400 and 500 base pairs, V8 at 300, 400, and 500 base pairs, and V9 at all lengths). This exclusion criteria was considered less than ideal, but necessary, as comparison of global and local tree topology metrics of trees of different sizes introduces considerable difficulty of interpretation of results. For each set of short reads, we calculated aforementioned global tree distance metrics, local cladal similarity metrics, as well as MAST sizes as compared to gold standard trees. We used beta regression with a logit link function to model global tree distance metrics (RF distance) and cladal similarity metric (shared sister pairs) as a function of short-read length (Length), variable region (VR), and the presence of guides (Group), including all main and two way interactions (betareg 3.2.1; *betareg(RF_Distance ∼ Length + VR + Group + VR*Group + VR*Length + Group*Length, link = “logit”)*. We evaluated the effect of guides using Wilcoxon Rank Sum Tests to determine whether the inclusion of guides significantly improved tree similarity scores. A Pearson correlation was used to determine if other global and local distances / similarities were correlated to one another.

Hypothesis 3 sought to determine if guide sequences could improve accuracy of phylogenetic inference when combining short-reads derived from two different but overlapping VRs. We simulated short-reads from the V4 or V3V4 region, and additionally constructed a mixed, overlapping dataset by randomly selecting simulated short reads without replacement from either group with equal probability. This resulted in three distinct short-read simulation groups – V4, V3V4, and a V4-V3V4 mixed group. Short-reads were either analyzed alone or combined with curated full-length guide sequences for phylogenetic inference (Figure 2b.S3 and 2c), and tree dissimilarity/similarity metrics were calculated as described previously. We evaluated the effect of guides using Wilcoxon Rank Sum Tests, and then correcting for multiple hypothesis testing using false discovery rate correction implemented with the *p.adjust(method = “fdr”)* function in the stats package (version 4.2.1).

Hypothesis 4 sought to determine if guide sequences could improve accuracy of combining short-reads derived from disjoint (i.e. non-overlapping) VRs. We simulated short-reads from the V1V2, V3V4, or V5V7 region, and additionally constructed a mixed, non-overlapping dataset by randomly selecting simulated short-reads without replacement from each group with equal probability. This resulted in four distinct short-read simulation groups. Short-reads were either analyzed alone or in combination with curated full-length guide sequences for phylogenetic inference (Figure 2b.S4 and 2c). We evaluated the effect of guides as before.

Hypothesis 5 sought to determine if guide sequences could improve accuracy of combining short-reads derived from a diverse mix of VRs – as might be the case for a typical meta-analysis. We simulated short-reads that are commonly found in experimentally derived microbial 16S studies (V1V2, V1V3, V3V4, V3V5, V4, and V4V5). As before we additionally created a mixed dataset by randomly selecting simulated short-reads without replacement from each group with equal probability. Short-reads were either analyzed alone or in combination with curated full-length guide sequences for phylogenetic inference (Figure 2c). We evaluated the effect of guides as before.

Hypothesis 6 sought to determine if the application of guides would show similar results when applying our method to experimental data. We analyzed raw 16S rDNA sequencing datasets generated from both Illumina short-read and PacBio long-read platforms using the same underlying sample set. A total of 146 human microbiome samples (saliva, oral biofilm, and fecal) were downloaded for the National Center for Biotechnology Information Sequence Read Archive (SRA accession project # PRJNA933120), comprising 26 Illumina and 120 PacBio runs. Dada2 (*version 1.30.0*) was used to remove primers, quality filter reads, exclude chimeras, and infer short- and long-read amplicon sequence variants (ASVs) in R (*version 4.3.3*) using custom scripting. Illumina short reads (*truncLen = c(275), maxEE = c(2), merge = FALSE*) and PacBio long reads (*minLen = 050, maxLen = 1700, maxEE = 4, maxN = 1, minQ =2*) were quality filtered, and then taxonomic annotations were assigned to all ASVs (SILVA; version 138.1), and any sequences not taxonomically annotated as Bacteria or Archaea were excluded from downstream analyses.

We considered the subset of long- and short-read ASVs for which a one-to-one pairing could be established – all other ASVs were discarded from downstream analysis. We considered trees built from long-read sequences to be closest to gold standard trees. As before we used short-read sequences alone, or combined with curated full-length guide sequences for phylogenetic inference. Guide sequences were identified by blasting short-read sequences to the SILVA database^30^ (*version 138.2 NR99 fasta file*) to identify those long-read sequences which matched short-read sequences by at least 99% identity, and had 100% coverage of the query sequence (*-task blastn -dust no – strand both -evalue 1e-6 -perc_identity 99 -qcov_hsp_perc 100*). We reasoned that this would provide the most similar suite of full-length guide sequences to our short-read inputs. Short-read tree similarity or dissimilarity was calculated as before, and the impact of guides was determined using the Wilcoxon Rank Sum test.

### Data visualization

The 16S model *E. coli* sequence was built using the webserver version of R2DT^55^, a method for predicting and visualizing RNA structures in standard layouts. Data were visualized by custom scrips in R (*version 4.2.1*) using ggplot2 (*version 3.5.1*).

### PhyloguidesR

We provide an R wrapper which enables easy use of our methods developed here, and an online tutorial for researchers to apply these tools to any set of input sequences and guides (https://github.com/HollyKArnold/phyloguidesR/).

### Artificial Intelligence (AI)

Our use of AI to prepare this manuscript was conducted in accordance with the polices of the American Society of Microbiology AI authorship guidelines at the time of writing last accessed on December 16^th^, 2025. We used AI primarily for copy editing to improve clarity, and readability. Specifically, we used GPT-4o version 2024.02.13 provided by OpenAI^56^, accessed from August 21^th^, 2024 to August 7^th^, 2025, GPT-5 version 2024.02.13 accessed from August 7^th^, 2025 to November 11^th^, 2025, and GPT 5.1 accessed from November 12^th^, 2025 to December 17^th^ 2025.

## Acknowledgements

I would like to thank Martha F. Witkop for editorial comments. Thank you Ian Humphreys for performing pilot simulations and editorial comments. National Science Foundation grant 1557192 and internal grants contributed funding towards this project.

## Author Contributions

H.K.A. conceptualized experiments, performed simulation experiments, analyzed data, visualized data, and wrote paper. E.L. prepared experimental data and wrote corresponding methods section. A.H. performed initial simulation pilot experiments. T.J.S. conceptualized experiments, provided edits on the paper, and oversaw the study. All authors read and approved final draft.

## Data Availability

The phyloguidesR package is available online at (https://github.com/HollyKArnold/phyloguidesR/).

## ABBREVIATIONS

VR: Variable Region
16S: 16S rDNA gene
RF: Robinson Foulds
JRF: Jaccard Robinson Foulds
MSD: Matching Split Distance
PID: Phylogenetic Information Distance
MCD: Mutual Clustering Distance
PD: Path Distance
QUAR: Quartet Distance

